# Water availability dynamics have long-term effects on mature stem structure in *Vitis vinifera*

**DOI:** 10.1101/265207

**Authors:** S. Munitz, Y. Netzer, Ilana Shtein, A. Schwartz

## Abstract

*Vitis vinifera* is a climbing vine with wide vessels and high hydraulic conductivity. There is a lack of data on the anatomical structure of the mature vine stem, and most current knowledge is based on first-year shoots. Moreover, the effect of drought stress on anatomical structure has been partly reported in shoots of *Vitis vinifera* but not in stems.

In current study two irrigation approaches were applied on *Vitis vinifera* Merlot vines: constant (low, medium and high irrigation) and dynamic (early/late season water deficit). The following parameters were measured: trunk diameter, annual ring width and area, vessel diameter, specific hydraulic conductivity and stem water potential.

High water availability early in the season (high irrigation and late deficit) resulted in vigorous vegetative growth (greater trunk diameter, ring width and area), wider vessels and increased specific hydraulic conductivity. The distribution of large xylem vessels was altered by drought stress, where high water availability early in the season caused a shift of the vessel population towards the wider frequency classes. Interestingly, the early deficit vines showed more negative water potential values late in the season compared to the low irrigation vines. This may imply an effect of anatomical structure on vine water status.

**Highlights:** Water availability early in the season determines vegetative growth and stem anatomical structure in mature *Vitis vinifera* vines.

## Introduction

The genus *Vitis* has been distinguished for its wide (Adkinson 1913; Pratt 1974) and long xylem vessels (Zimmermann and Jeje 1981; Ewers et al. 1990; Jacobsen et al. 2012), which characterize lianas. Such hydraulic architecture makes ***Vitis vinifera*** an excellent model of water flow in plants according to the “unit pipe model” as described by Tyree & Ewers (1991), since *Vitis* vessels are relatively optimal pipes. Since vessel length and diameter are correlated with stem diameter, mature vine trunks tend to have wider and longer vessels compared to young stems/shoots (Ewers and Fisher 1989; Jacobsen et al. 2012; Hacke 2015). Moreover, since vessels in *Vitis* are completely inactivated after 4 to 7 years (Tibbetts and Ewers 2000), in mature vine trunks (older than 7 years) only the secondary xylem is functional, while the primary xylem created in its first year is nonfunctional. This fact is crucial, since in contrast to the scalariform arrangement of intervessel bordered pits of secondary xylem vessel elements, the primary xylem vessel elements have partial secondary wall thickenings, making them much more vulnerable to cavitation (Pratt 1974; Choat et al. 2005; Sun et al. 2006; Brodersen et al. 2011; Craig R Brodersen et al. 2013; Rolland et al. 2015; Hochberg, Herrera, et al. 2016). Indeed, visualization techniques (microCT / MRI) have shown that in one-year-old stems of *Vitis* the spread of embolism proceeds from the pith towards the cambium through the primary xylem (Choat et al. 2010; Craig R Brodersen et al. 2013; Craig Robert Brodersen et al. 2013; Knipfer et al. 2015; Hochberg, Albuquerque, et al. 2016). An additional important anatomical feature of mature vine trunks is the ratio between the area of the pith and the xylem, which decreases with stem maturation (Sun et al. 2006). All of the above suggest that the xylem architecture of the mature trunk differs from that of a young shoot, in a way that makes it less vulnerable to cavitation.

Despite those significant differences, there is a scarcity of information about the anatomical structure of mature vine trunks, while current reported anatomical information on vines is based mainly on analysis of one-year-old stems (Schultz and Matthews 1993; Lovisolo et al. 1998; Schubert et al. 1999; Sun et al. 2006; Brodersen et al. 2011; Chatelet et al. 2011; Santarosa et al. 2016). In one-year-old stems, hydraulic structure is reported to vary among *Vitis* cultivars and rootstocks (Chouzouri and Schultz 2005; Chatelet et al. 2011; Gerzon et al. 2015; Hochberg et al. 2015; Santarosa et al. 2016; Shtein et al. 2016), and to be affected by environmental parameters (Schubert et al. 1999). Only a few studies analyzing anatomical features of xylem in mature *Vitis* trunks have been published (Zimmermann and Jeje 1981; Ewers et al. 1990; Tibbetts and Ewers 2000; Shtein et al. 2016).

Another interesting feature of *Vitis* xylem is the bimodal distribution pattern of vessel diameters, meaning two distinct vessel size groups – wide and narrow (Carlquist 1985; Ewers et al. 1990; Wheeler and LaPasha 1994; Shtein et al. 2016). Wide diameter vessels are considered to be more hydraulically efficient, and tend to be more vulnerable to embolism within the same species (Sperry and Tyree, 1988; Lo Gullo and Salleo, 1991; Hargrave et al., 1994; Cai and Tyree, 2010; Christman et al., 2012; Scoffoni et al., 2016). The accepted “air-seeding” theory suggests that the increased vulnerability to embolism of wide vessels is linked to their enlarged total area of intervessel pits. A wide pit area raises the average size of the “rare” largest pore, consequently increasing the risk of air seeding (Choat et al. 2003; Wheeler et al. 2005; Jansen et al. 2009; Cai and Tyree 2010).

Most cultivated vineyards worldwide are located in semi-arid and arid regions where drought stress is prevalent (Chaves et al. 2007). Nevertheless, compared to other woody plants, grapevines are often described as relatively vulnerable to drought stress (Choat et al. 2010; Zufferey et al. 2011; Jacobsen and Pratt 2012; Hacke 2015). It has been reported that drought stress induces embolism and a loss of hydraulic function (Schultz and Matthews 1988; Hargrave et al. 1994; Lovisolo et al. 1998; Choat et al. 2010; Brodersen et al. 2014). Drought stress also negatively affects vegetative growth and pruning weight of vines (Matthews 1987; Intrigliolo and Castel 2010; Shellie and Bowen 2014; Munitz et al. 2016). There is a lack of available information on the effects of drought stress on hydraulic conductivity resulting from modifications to *Vitis* xylem anatomy, as only a few studies have examined this subject (Lovisolo et al. 1998; Hochberg et al. 2015). The current study focuses on the long-term anatomical acclimation to drought stress of mature *Vitis vinifera* Merlot vines.

## Materials and Methods

### Plant material and experimental design

This study was carried out in a 100-ha commercial vineyard located in the Judean Plain, Israel (31^0^49′N, 34^0^53’E, elevation 124 m). This region has a semiarid climate with predominantly winter rainfall (average 463 mm year^−1^) and high evapotranspiration (average 1512 mm year^−1^). The vineyard was planted in 1998 with Vitis *vinifera* L cv. ‘Merlot’ grafted to 140 Ruggeri, and trained onto a two-wire vertical trellis. Row direction was North/South with a slight tendency to the West, and vine and row spacing were 1.5 m and 3 m respectively (2222 vines ha ^−1^). The soil was loam (48% sand, 29% silt and 23% clay, field capacity 28 % vol., wilting point 14 % vol.). Pest management and fertilization in the vineyard were applied according to standard local agricultural practice.

The experimental design was a complete randomized block design with five irrigation treatments each replicated four times. Each block comprised three rows (one data and two border rows). Each plot comprised 16 vines per line, with the outer two vines at each end being buffer vines and the inner 12 vines being measurement vines (a total of 240 measurement vines, i.e. 12 vines × 5 treatments × 4 replicates).

### Irrigation treatments

During 2009-2012, five irrigation treatments representing different levels of deficit irrigation were applied as percentages of crop evapotranspiration (ET_c_). Crop evapotranspiration was calculated by multiplying reference evapotranspiration (ET_o_) by the crop coefficient (K_c_), i.e. ET_c_ = ET_o_× K_c_. ET_o_ was calculated using data obtained from the adjacent meteorological station, and K_c_ was calculated according to Netzer et *al* (2009) following nondestructive measurements of leaf area index. For more details about the irrigation method see Munitz and Netzer (2016). Irrigation treatments followed two different strategies: static irrigation and dynamic (seasonally changing) irrigation. Dynamic treatments involved alternation of the percentage of ET_c_ along the growing season according to phenological stages (stage I, stage II, stage III) as defined by Kennedy (2002): stage I – from bloom to bunch closure, stage II – from bunch closure to veraison (color change to red) and stage III – from veraison to harvest. Dynamic irrigation treatments were: early deficit (0, 20, 50% of ET_c_) and late deficit (50, 20, 20% of ET_c_). Static irrigation treatments were: low irrigation (20% of ET_c_), medium irrigation (35% of ET_c_) and high irrigation (50% of ET_c_).

### Stem water potential (ψ_s_)

Stem water potential (ψ_s_) was measured using a pressure chamber (Arimad 2, Kfar Charuv, Israel). Three sunlit, mature, fully-expanded leaves from each plot (12 leaves per treatment) were bagged 2 h prior to measurement in plastic bags covered with aluminum foil. The time elapsing between leaf excision and chamber pressurization was less than 15 s. The measurements were conducted one day before irrigation was applied.

### Trunk diameter

Measurements were performed monthly with a digital caliper (075430, Signet, Taiwan) on 48 vines per treatment (12 vines per plot × 4 replicates). In order to obtain consistent data, all measured vines (240) were marked 30 cm above ground with colored tape, and all measurements were taken at this point.

### Anatomical sampling

At the end of the experiment (December 2012) xylem cores from representative vines were sampled 50 cm above ground with an increment borer (5.15 mm Core 3-Thread 8”, Haglof, Sweden). Twelve cores were sampled from each treatment (3 cores per plot × 4 replicates, 60 cores total). Trunk diameter (*D*; mm) at the drilling location was recorded. Cores were placed in sterilized water and stored at 4° C until cross sectioned with a sliding microtome (NR17800, Reichert, Austria) at a thickness of 90 μm. In order to increase visual contrast, cross sections were stained for 60 s in Reactif Genevois solution (FAHN 1954), then flushed with distilled water. Photographs of stained cross sections were obtained using a stereo microscope (Olympus SZ2-ILST) coupled with a digital camera (Olympus LC20) equipped with image acquisition software (LCmicro 5.1, Olympus, Tokyo, Japan) at ×20 magnification.

### Image analysis

Analysis of cross sections was performed by separately quantifying various parameters in the visible field for each of four recent growth rings (2009-2012) using ImageJ software (Rasband, W.S., ImageJ, U.S. National Institutes of Health, Bethesda, Maryland, USA, http://imagej.nih.gov/ij/, 1997-2016). The abbreviations for structural parameters were taken from Scholz *et al.* (2013). The following anatomical parameters were measured (Fig. 1): annual ring width (*W*_r_; μm), bark width (*W_b_;* μm), xylem radius (*r_x_*; μm) and inner xylem radius (*r;_i_* μm). Vessel lumen area (*A*_v_; μm) and number of vessels (*n*) were measured using the ‘analyze particles’ tool (see Fig. 2 for explanation); analyzed area (A; mm^2^) was also measured. A total of 12,177 vessels were measured and used for subsequent hydraulic conductivity calculations. The detailed calculations for trunk and vessel parameters are presented in Table 1.

**Figure 1.**
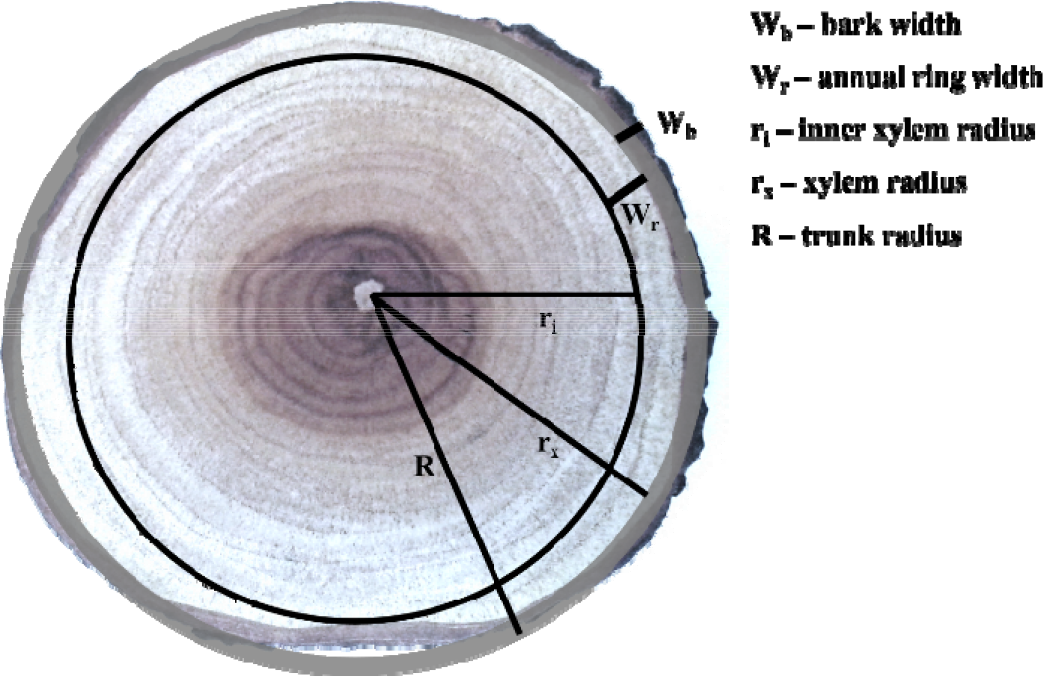
Schematic cross section diagram with abbreviations used for trunk parameter calculations.

**Figure 2.**
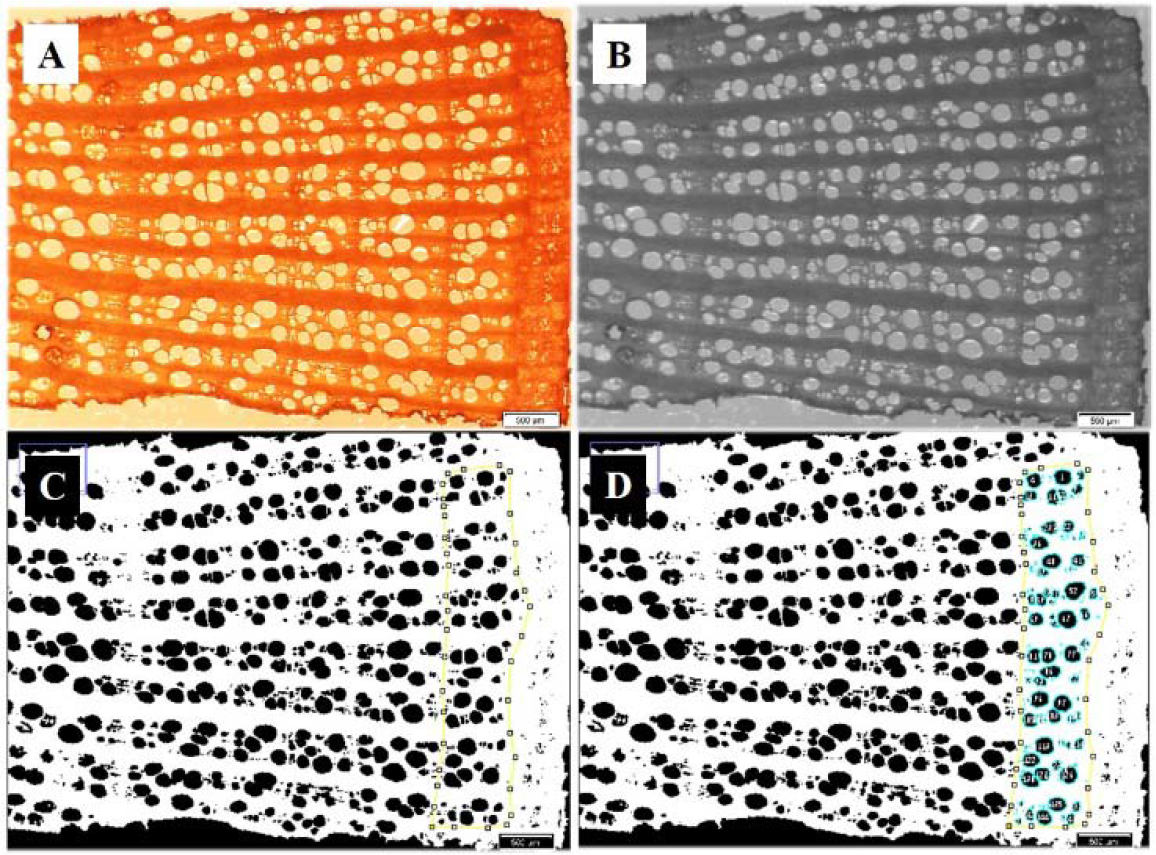
Stem cross section of vitis vinifera cv. Merlot. (A) Cross section image as acquired by stereo microscope. (B) Image processing to 8-bit. (C) Image conversion to black and white (binarization) and selection of annual ring to be analyzed (yellow line). (D) Measurement of vessel area.

**Table 1.** Calculations used for trunk and vessel parameters.

| Parameter | Abbreviation | Unit | Formula |
| --- | --- | --- | --- |
| Vessel diameter | $d$ | $\mu\text{m}$ | $d = (4 * A_v / \pi)^{0.5}$ |
| Vessel density | $V_D$ | $\text{mm}^{-2}$ | $V_D = n/A$ |
| Trunk radius | $R$ | $\mu\text{m}$ | $R = D/2$ |
| Xylem radius | $r_x$ | $\mu\text{m}$ | $r_x = R - W_b$ |
| Annual ring area | $A_r$ | $\text{mm}^2$ | $A_r = \pi * (r_x)^2 - \pi * (r_i)^2$ |

### Theoretical specific hydraulic conductivity (K_s_) calculations

The theoretical specific hydraulic conductivity (*K_s_*; kg m^−1^ MPa^−1^ s^−1^) was calculated using the modified Hagen–Poisseuille’s equation (Tyree and Ewers 1991):

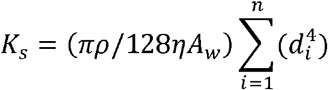

where *K_s_* is the specific hydraulic conductivity, *p* is the density of the fluid in kg m^−3^ (assumed to be 1000 kg m^−3^), η is the dynamic viscosity of the fluid in MPa s^−1^ (assumed to be 1×^−9^ MPa s^−1^), *A_w_* is the area (m^2^) of the xylem cross section analyzed, *d* is the diameter (m) of the i^th^ vessel and n is the total number of vessels in the measured area.

Hydraulic conductivity per annual ring *(K_ar_*, kg m^1^ MPa^−1^ s^−1^) was calculated by multiplying the theoretical xylem specific hydraulic conductivity (*K_s_*) by annual growth ring area (*A_r_*s, m^2^).

Frequency classes for calculation of total vessel number and total conductivity were established at intervals of 20 µm (Fig. 3). For calculation of vessel density and vessel average diameter, vessels were separated into two size categories (>100 μm, ≤ 100 μm), since *Vitis* has a bimodal distribution of xylem vessels (Fig. 3A).

**Figure 3.**
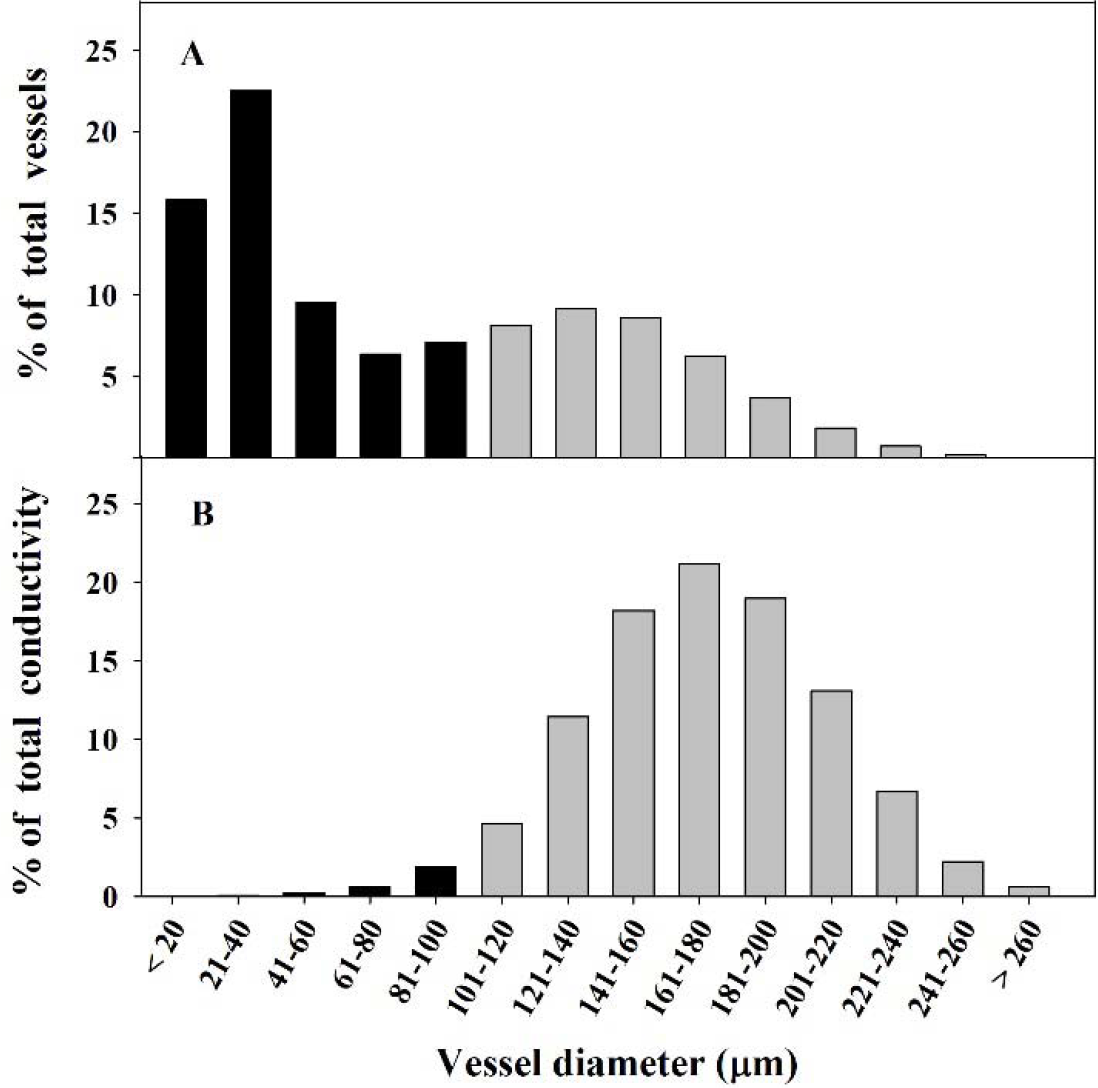
(A) Distribution of stem xylem vessels according to diameter classes (μm) in Merlot vines of all irrigation treatments. (B) Distribution of hydraulic conductivity according to diameter classes (μm) in Merlot vines of all irrigation treatments. Vessel classes were divided into two size categories: ≤100 μm (black bars) and ≤100 μm (gray bars). Data represent vessels from all irrigation treatments during 2009-2012, n = 12177 vessels.

### Statistical analysis

The software program JMP7 (SAS Institute, Cary, NC) was used for all statistical procedures. Data were analyzed via analysis of variance (ANOVA), and means were separated according to the least significant difference (LSD) at p ≤ 0.05 using the Tukey-Kramer test.

## Results

### Stem vessel distribution

The distribution of stem xylem vessels in all irrigation treatments showed a classic bi-modal pattern. The small vessels (< 100 μm) constituted the majority (61.3%) of total vessel number, while the large vessels comprised only 38.7% of total vessels (Fig. 3A). In contrast, the theoretical hydraulic conductivity showed a reverse distribution, where the large vessels contributed 97.2% of total conductivity (Fig. 3B), whereas the contribution of the more abundant small vessels was negligible.

### Seasonal changes in trunk diameter

As an integrative indicator of vegetative growth, seasonal changes in trunk diameter were monitored monthly during 2011-2012 (following two years of differential irrigation application after 11 years of identical irrigation). Seasonal trends of trunk diameter development were similar in both years in all irrigation treatments (Fig. 4); an increase in trunk diameter began two weeks after bud break and continued until stage II (mid-June), then remained stable until the next season. In 2011, a decrease in trunk diameter was apparent during stage III, in all irrigation treatments. Vines in different irrigation treatments showed significant differences in trunk diameter throughout the entire experimental period (Fig. 4). Among static irrigation vines, trunk diameter was positively correlated with applied water amounts, even though the trunk of the medium irrigation vines was slightly narrower than expected (Fig. 4A). In the dynamic irrigation treatments, the early deficit vines exhibited the narrowest trunk diameter of all vines in all irrigation treatments during the entire measuring period. The late deficit vines had an intermediate trunk diameter throughout the measuring period (Fig. 4B).

**Figure 4.**
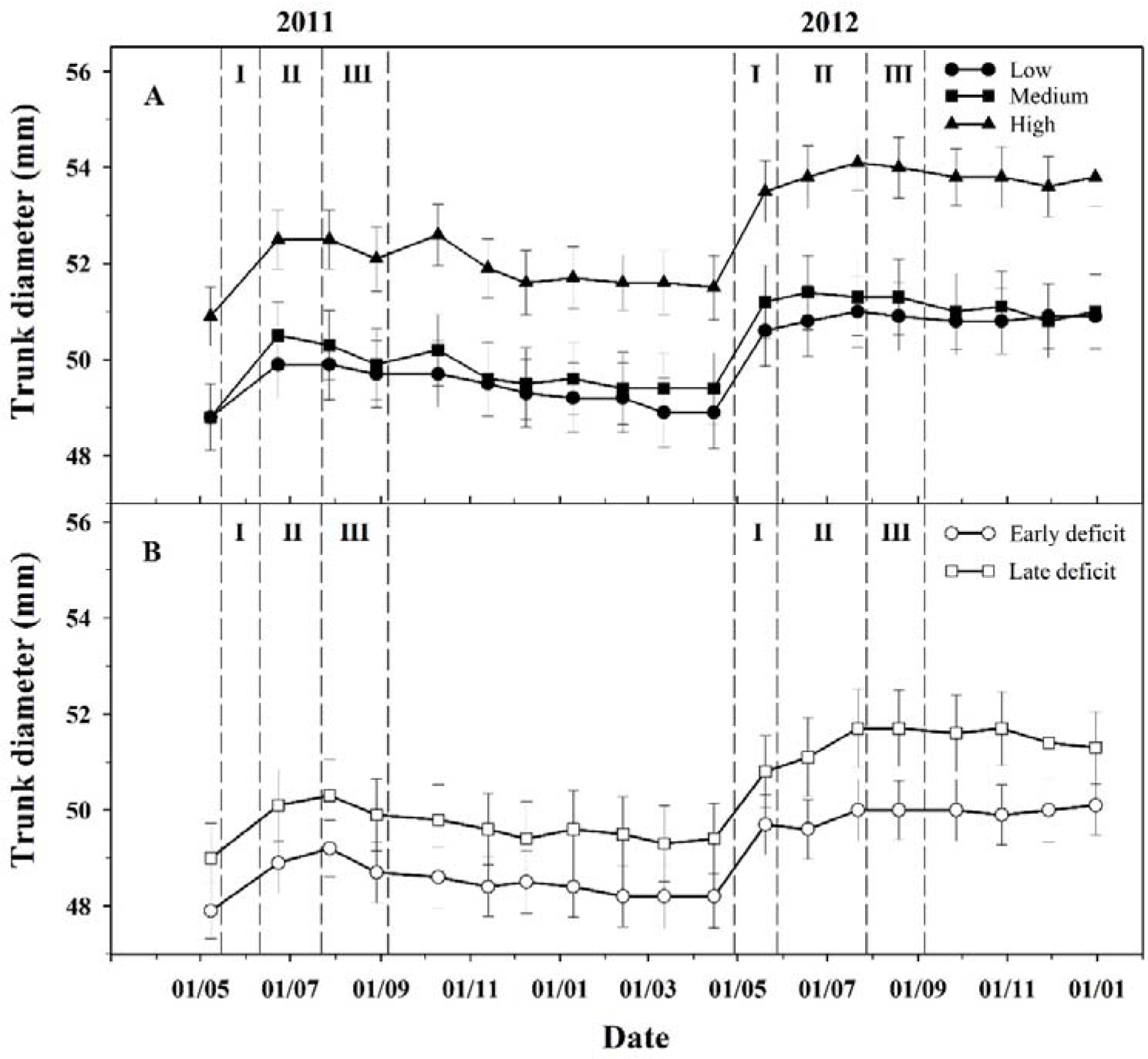
(A) Biennial pattern of trunk diameter development of Merlot vines exposed to low irrigation (closed circles), medium irrigation (closed squares) and high irrigation (closed triangles), in 2011 and 2012. (B) Biennial pattern of trunk diameter development of Merlot vines exposed to early deficit (open circles) and late deficit (open squares), in 2011 and 2012. Each point is the mean of 48 vines (12 vines per replicate). The vertical bars denote one standard error.

### Structural parameters

Vines subjected to the static irrigation treatments differed significantly in their annual ring width, with a positive effect of applied water amounts on ring width (Table 2). Within the dynamic irrigation treatments, the late deficit vines had the widest annual ring width (901.5 μm), very similar to the width of the high irrigation vines (Table 2). Surprisingly, the early deficit vines had the narrowest annual ring width (686 μm) even compared to the low irrigation vines (719 μm). The general trend of annual ring area resembled the trend of annual ring width. Ring area was positively affected by increasing water amounts in static treatment vines. In dynamic irrigation treatments, early deficit vines had the smallest annual ring area of the five treatments, while the late deficit vines had a relatively large ring area, intermediate between that of the high and medium irrigation vines (Table 2).

**Table 2.**
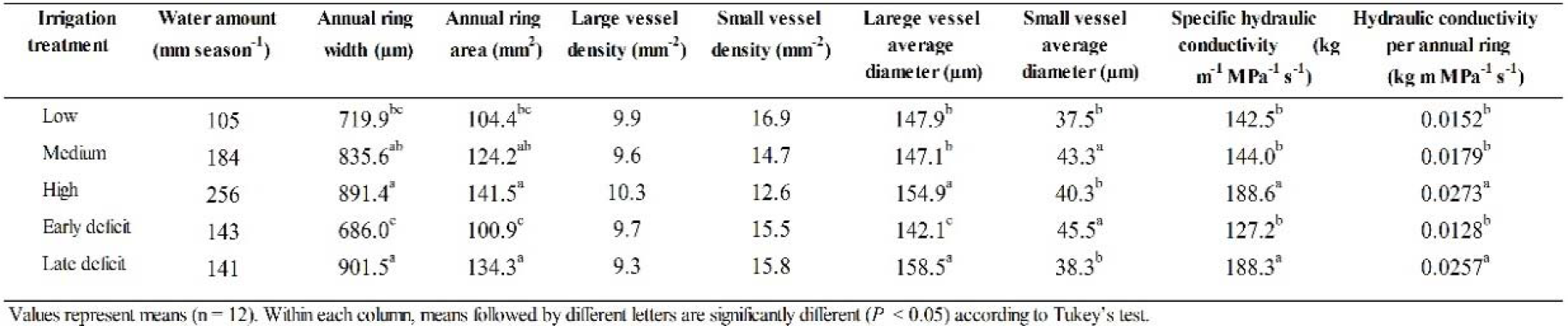
Water amounts, anatomic and hydraulic parameters. “Hulda’ Merlot vineyard, 2009 - 2012.

The density of ‘small’ (≤100μm) and ‘large’ (>100μm) vessels was not significantly affected by irrigation treatments, although a reduction in the density of ‘small’ vessels in high irrigation vines was observed. The ‘large’ (>100μm) vessel diameter of the high irrigation vines was significantly wider than that of the low and medium irrigation vines. In the dynamic irrigation treatments, early deficit vines had significantly narrower ‘large’ vessels than all vines in all other irrigation treatments, while the late deficit vines had the wideset (Table 1). The trend for ‘small’ (≤100μm) vessel diameter was less clear, with early deficit and medium irrigation vines exhibiting significantly wider vessels than all other vines. The trend in specific hydraulic conductivity was similar to that of ‘large’ (>100μm) vessel diameter, where the high irrigation and late deficit vines had significantly higher hydraulic conductivity (Table 1) than vines in other treatments. Hydraulic conductivity per annual ring increased significantly with increasing water amounts in the static irrigation treatments, whereas in dynamic treatments the early deficit vines had the lowest conductivity, while the late deficit vines had high conductivity (slightly lower than the high irrigation vines). The relationship between hydraulic conductivity per annual ring and seasonal water amount (Fig. 6a) was weak and non-significant (R^2^ = 0.21), while the relationship between hydraulic conductivity per annual ring and water amounts applied during stage I (bloom to bunch closure, Fig. 5b) was stronger and significant (R^2^ = 0.60, p < 0.001).

**Figure 5.**
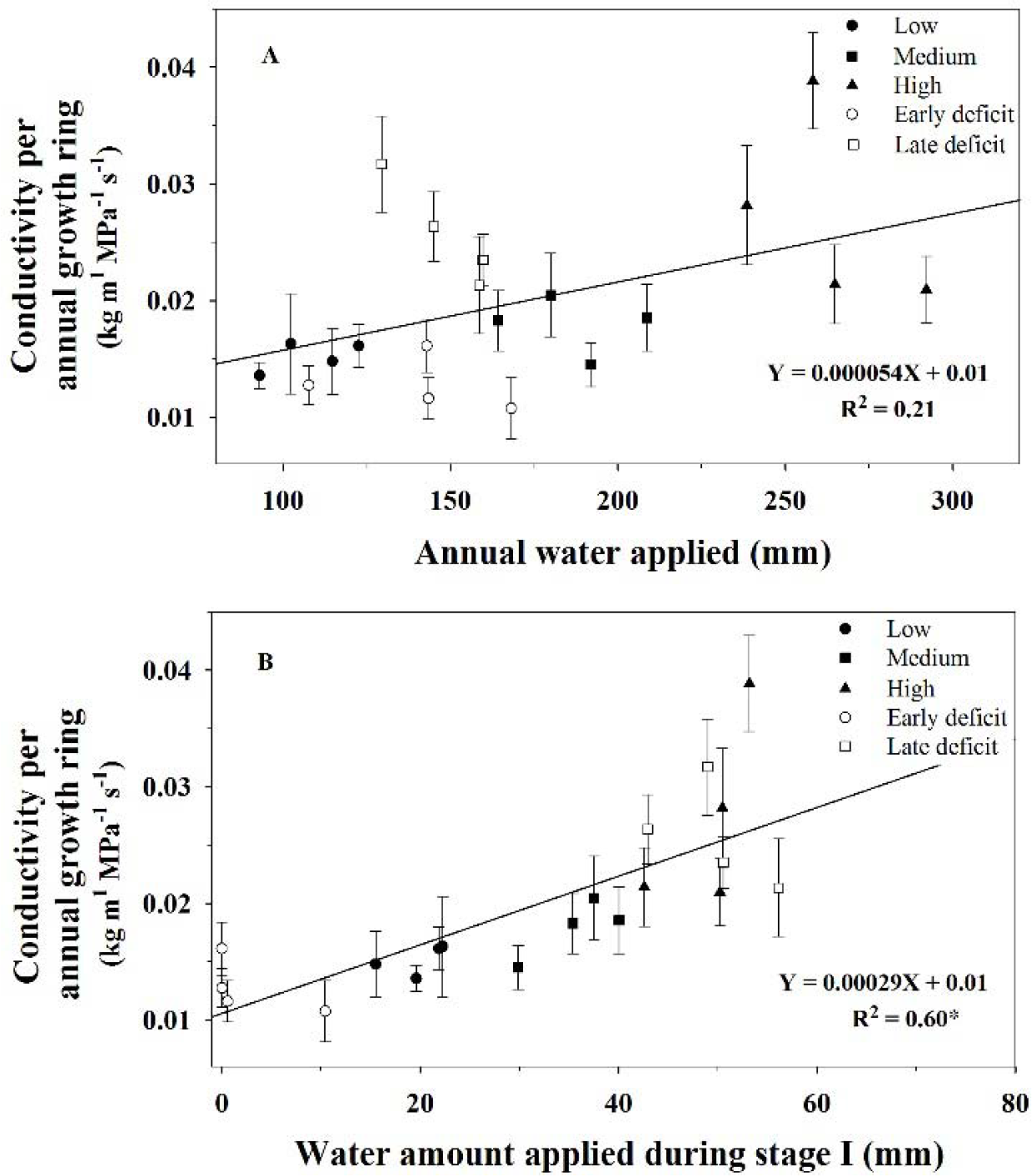
(A) Relationship between annual water applied and hydraulic conductivity per annual growth ring in Merlot vines of all irrigation treatments. (B) Relationship between water amount applied during stage I (bloom to bunch closure) and hydraulic conductivity per annual growth ring in Merlot vines of all irrigation treatments. Each point is the mean of 12 vines (3 vines per replicate). The vertical bars denote one standard error. *Significant at P < 0.001.

**Figure 6.**
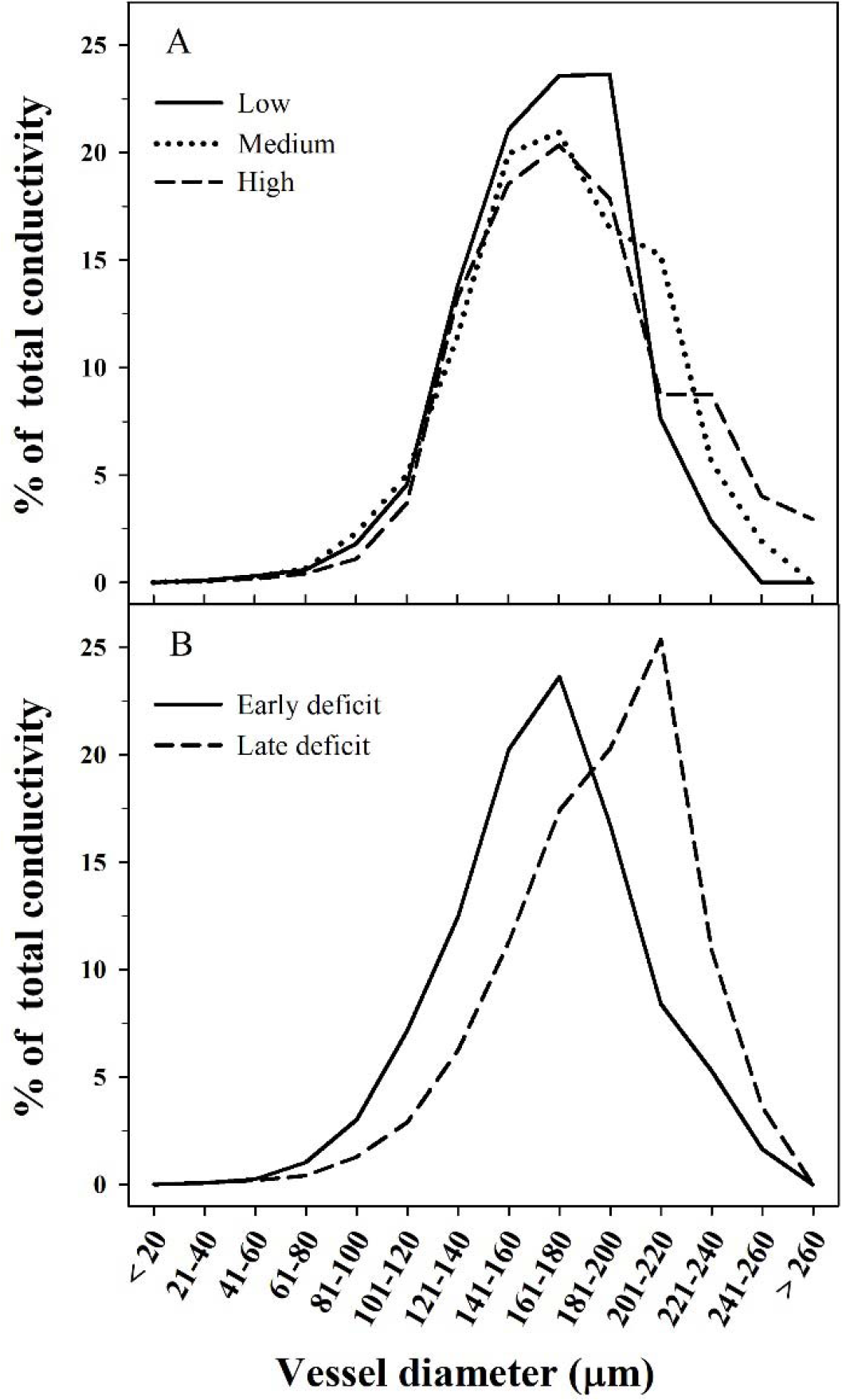
(A) Distribution of hydraulic conductivity according to diameter classes (μm) in Merlot vines exposed to low irrigation (solid line), medium irrigation (dotted line) and high irrigation (dashed line). (B) Distribution of hydraulic conductivity according to diameter classes (ym) in Merlot vines exposed to early deficit (solid line) and late deficit (dashed line). Data represent vessels from all analyzed years (2009-2012), n = 12177 vessels.

### Stem hydraulic conductivity distribution among vessel size classes

Specific stem hydraulic conductivity was calculated separately for each vessel size class (Fig 6). Among static irrigation treatments, the conductivity distribution was similar, but several differences among treatments were observed. In low irrigation vines a higher percentage of the calculated hydraulic conductivity was derived from narrow vessel classes, in the medium irrigation vines – from wide vessel classes, and in the high irrigation vines – from the widest vessel classes (Fig 6a). In the dynamic irrigation treatments, the early deficit vines had a higher amount of hydraulic conductivity derived from narrower frequency classes, while the hydraulic conductivity in late deficit vines shifted towards the wide frequency classes (Fig 6b).

### Stem water potential

Significant differences in water potential (ψ_s_) between vines in different irrigation treatments were observed during stages II and III (Fig. 7). The daily trend of ψ_s_ was similar among treatments, with a steep decrease in ψ_s_ values being recorded throughout the morning, followed by stabilization and improvement in Ts during the afternoon (Fig. 7). In stage II an improvement in Ts values during the afternoon was apparent only in the high and medium irrigation vines (Fig. 7B). On both measuring days, the high irrigation vines had the highest Ts values during the course of the day, the medium irrigation vines were at an intermediate level and the low irrigation vines had the lowest ψ_s_. In the dynamic irrigation treatments, the early deficit vines had low ψ_s_ values during stage II (lower than the low irrigation vines) and intermediate values during stage III (resembling those of the medium irrigation vines). The late deficit vines had low ψ_s_ values on both dates, even compared to the values of the low irrigation vines (Fig. 7).

**Figure 7.**
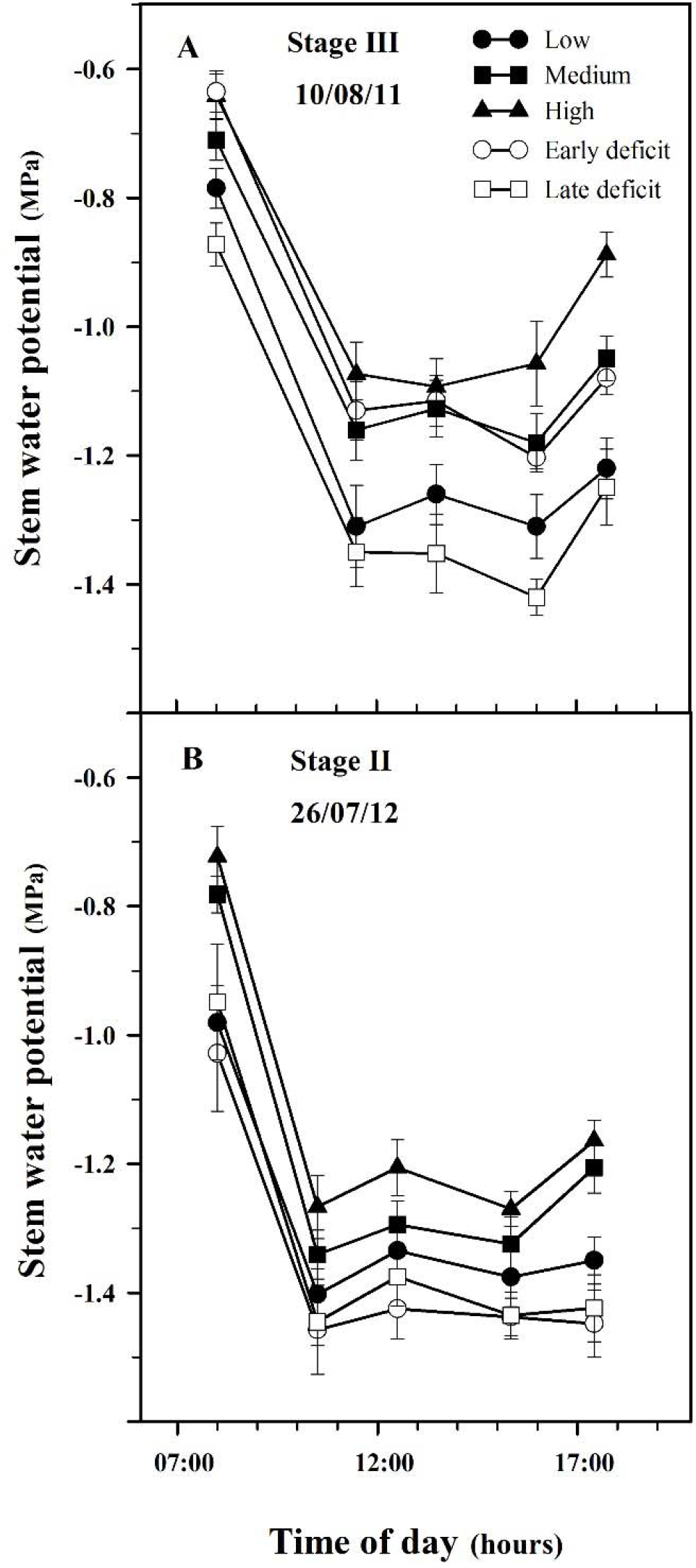
(A) Daily pattern of stem water potential of Merlot vines exposed to low irrigation (closed circles), medium irrigation (closed squares), high irrigation (closed triangles), early deficit (open circles) and late deficit (open squares), on 10/08/11. (B) Daily pattern of stem water potential of Merlot vines exposed to low irrigation (closed circles), medium irrigation (closed squares), high irrigation (closed triangles), early deficit (open circles) and late deficit (open squares), on 26/07/12. Each point is the mean of 12 leaves (3 vines per replicate). The bars denote one standard error.

## Discussion

### Vegetative growth

Vegetative growth can serve as a good indicator of water availability (Tyree and Ewers 1991; Munitz et al. 2016). In the present study we continuously monitored trunk growth. In all irrigation treatments, an increase in trunk diameter occurred mainly early in the season (from two weeks before blooming until bunch closure) (Fig. 4). Early season trunk growth has been previously reported in Merlot (Ton and Kopyt 2004) and in other *Vitis vinifera* cultivars (Myburgh 1996; Ton and Kopyt 2004; Intrigliolo and Castel 2007; Montoro et al. 2011; Papi and Storchi 2012; Edwards and Clingeleffer 2013).

The high irrigation and late deficit vines, which received higher water amounts early in the season, exhibited the widest trunk diameter of all vines. Similarly, annual ring width and area, which represent annual vegetative growth, were also positively correlated with high irrigation early in the season (Stage I). The high irrigation and late deficit vines had significantly wider ring width and area in comparison to the low irrigation and early deficit vines (Table 2). The dominance of early season vegetative growth in *Vitis vinifera* can be explained by the fact that cambial activity to produce new vascular elements takes place mainly during the early stage of the growing season (until 20 days after bunch closure, Bernstein and Fahn, 1960).

### Hydraulic structure

A significant increase (9 - 11%) in the large vessel diameter was recorded in the high irrigation and late deficit vines (Table 2), even though no significant difference in the density of the large vessels was found. This implies that water availability early in the season affects large vessel diameter rather than vessel density. The same effect of drought stress reducing average vessel diameter has been reported in *Vitis vinifera* shoots (Lovisolo et al. 1998) and petioles (Hochberg et al. 2015). Interestingly, in small vessels the opposite was found, where the vessels of the early deficit vines, which received minimal water early in the season, exhibited the widest diameter (Table 2). Although the small vessels comprised the majority of the total vessels (61%), they contributed only a negligible 3% of total hydraulic conductivity (Fig. 3). A similar phenomenon of dominance of wide diameter vessels with respect to total hydraulic conductivity has also been recorded in other species (Hargrave et al. 1994; Tibbetts and Ewers 2000).

From inspection of the hydraulic conductivity distribution, it can be deduced that high water amounts early in the season result in a higher percentage of hydraulic conductivity derived from wider frequency classes (Fig. 6). This phenomenon is more pronounced in late deficit vines, possibly due to their dynamic irrigation treatment. It is well known that in *Vitis*, as in many other lianas, wide vessels are formed at the beginning of the growing period and narrow vessels are formed at the end (extreme diffuse-porous), causing a bi-model distribution of vessels (Pratt 1974; Kozlowski 1983; Ewers et al. 1990; Wheeler and LaPasha 1994). Applying high irrigation amounts during the formation of wide vessels and reducing water allocation during the formation of narrow vessels (late deficit), should result in larger wide vessels and smaller narrow vessels. This will lead to an increased proportion of hydraulic conductivity being derived from wide vessels. As a result, late deficit vines are expected to be more susceptible to embolism formation in comparison to low irrigation vines.

Specific hydraulic conductivity (*K_s_*) represents the conducting efficiency of the xylem tissue (Tyree and Ewers 1991). The high irrigation and late deficit vines had significantly increased *K_s_* (24 – 33%) compared to other vines (Table 2). This means that limited water availability early in the growth season (bloom to bunch closure) results in decreased *K_s_*. Since there is a good correlation between measured and calculated hydraulic conductivity (Salleo et al. 1985; Hargrave et al. 1994; Lovisolo et al. 1998; Nolf et al. 2017) our theoretical *K_s_* results may indicate an actual change in the hydraulic conductivity of vines in response to drought stress.

Hydraulic conductivity per annual growth ring (*K_ar_*) is an integrated parameter combining both hydraulic conductivity (*K_s_*) and vegetative growth (trunk diameter and ring width). As with ring area and hydraulic conductivity, the high irrigation and late deficit vines had significantly increased hydraulic conductivity per annual ring (*K_ar_*, Table 2). Interestingly, the high irrigation vines had 9% higher *K_ar_* (not significant) than the late deficit vines. This difference can be attributed to the reduction of water amounts in late deficit vines during bunch closure, while cambial activity in the stem persisted for a further 20 days (Bernstein and Fahn 1960).

Reinforcement of the notion that most of the vegetative growth and xylem development occur early in the season can be found in the correlation between the amounts of applied water and hydraulic conductivity per annual ring, *K_ar_* (Fig. 5). While annual water applied had a weak and non-significant correlation with *K_ar_* (R^2^ = 0.21), water amount applied during stage I was strongly and significantly correlated to *K_ar_* (R^2^ = 0.6, P < 0.001).

### Water potential

Stem water potential is known to be a sensitive indicator of vine water status (Choné et al. 2001; Munitz et al. 2016). Indeed, on both measuring days static irrigation vines differed significantly in their stem water potentials, according to the water amounts applied, throughout the entire day. Those differences in stem water potential were present from bunch closure until harvest (Munitz et al. 2016). Interestingly, late deficit vines were more drought stressed (more negative stem water potential) than low irrigation vines on both measuring days, even though they received on average 31% more water prior to the measuring days. Those differences in stem water potential cannot be attributed to a broader canopy, since the late deficit and low irrigation vines had similar leaf area on measuring days (Munitz et al. 2016). Water potentials of −1.4 MPa, recorded on both measuring days, have been reported to cause a 30 – 80 % decrease in hydraulic conductivity of *Vitis vinifera* shoots(Alsina et al. 2007; Choat et al. 2010; Jacobsen and Pratt 2012).

The increased drought stress in late deficit vines can be explained by a greater hydraulic conductivity loss. As stated before, wide vessels are more susceptible to embolism formation. Late deficit vines, with a higher percentage contribution of wide vessels to total hydraulic conductivity, are expected to experience greater hydraulic loss at a similar water potential. Increased hydraulic loss will, in turn, lead to increased drought stress e.g. more negative water potential (Tyree et al. 1991). Another explanation for the increased drought susceptibility of late deficit vines compared with low irrigation vines, can be found by examining their Carlquist□s “vulnerability index” (1977). The index is calculated by dividing average vessel diameter by vessel density, where a lower value is interpreted as greater redundancy of vessels and improved capability of withstanding drought stress. The large vessels of the late deficit vines have an index of 17.0 while low irrigation vines have an index of 14.9.

### Structural parameters as compared to values from literature

Generally, our results seem to be in agreement with values presented in the literature. The annual increase in trunk diameter of mature vines measured in this study (1.5 – 2.5 mm, Fig. 4), is similar to that reported for a number of *Vitis vinifera* cultivars (0.5 – 3.5 mm) (Bernstein and Fahn 1960; Myburgh 1996; Ton and Kopyt 2004). Similarly, the range of annual ring width measured in this study (720 – 901 μm) is consistent with values reported for mature *Vitis vinifera* cultivars (100 – 1300 μm) (Perold 1927; Bernstein and Fahn 1960), and the range of annual ring area found in this study (104 – 134 mm^2^) resembles that of Cabernet Sauvignon vines (120 – 240 mm ^2^, Shtein *et al.*, 2016).

The average diameter of large vessels measured in this study (147 – 158 μm) is considerably smaller than that reported for *Vitis vinifera* cv. Cabernet Sauvignon. While in the current study water availability of Merlot vines in different irrigation regimes triggered a maximal difference of 11 μm in vessel diameter (Table 2), the difference between the two cultivars stands at more than 50 µm. This implies an inherent genetic distinction in stem xylem anatomy between *Vitis vinifera* cultivars, as reported previously for petioles and shoots (Chouzouri and Schultz 2005; Chatelet et al. 2011; Tombesi et al. 2014; Gerzon et al. 2015; Hochberg et al. 2015; Santarosa et al. 2016). Typical values of specific hydraulic conductivity of liana stems (Milburn 1979) are 65 – 349 kg m^−1^ MPa^−1^ s^−1^; our *K_s_* results (142 – 188 kg m^−1^ MPa^−1^ s^−1^) are in the middle of this range. Values of *K_ar_* reported for *Vitis vinifera* cv. Cabernet Sauvignon (0.07 – 0.11, kg m^1^ MPa^−1^ s^−1^) are considerably higher than those calculated in this study, due to the wider vessels and ring area measured in Cabernet Sauvignon vines.

## Conclusions

In the current study, we conducted a comprehensive structural analysis of the mature stem xylem of *Vitis vinifera*, combined with physiological measurements. The stem comprises the perennial part of the deciduous vine and its anatomical structure constitutes the long term “memory” of the vine, yet very little information about the anatomical features of mature *Vitis* stems is available. One example of this “memory” is the fact that the current year’s vine canopy develops while consuming water conducted through vessels differentiated in previous years. Most of the vine’s canopy develops in the early stages of the growing season, about 60 days from budbreak (Ben-Asher et al. 2006; Intrigliolo et al. 2008; Romero et al. 2010; Munitz et al. 2016). Cambial tissue begins its activity about two weeks after budbreak (Bernstein and Fahn 1960). Vessels become hydraulically active about four weeks after their initial differentiation is initiated (Halis et al. 2012; Hacke 2015): two weeks for differentiation, expansion and formation of secondary walls and at least two additional weeks for the creation of perforation plates by autolysis of axial cell walls. Practically this means that current year vessels are functional no less than 54 days after budbreak, when canopy development has almost ceased.

There is a lack of information about the effects of drought stress on *Vitis* xylem structure (Lovisolo et al. 1998), especially in mature stems. Our research can contribute to the understanding of mature stem xylem structure and how it is affected by drought stress in the long term.

## Acknowledgments

This study was partly sponsored by the Israeli Wine Grape Council. The authors thank Michal Akerman and Dan Aluf for their assistance in setting up this study, and the dedicated growers of Kibbutz Hulda: Silvio Feldman, Robbie Handel and Yigal Gad. We particularly thank Yechezkel Harroch and Elazar Quinn for assisting in the field and to Ami Charitan and Ziv Charit from Netafim for donating the irrigation system. We thank the agronomists of Barkan Winery, Elad Gutman and Yakov Cohen-Achdut for their collaboration. We also thank Eran Harkabi from the Israel Ministry of Agriculture and Rural Development for his assistance.References.

